# Coherent coding of spatial position mediated by theta oscillations in hippocampus and prefrontal cortex

**DOI:** 10.1101/520585

**Authors:** Mark C. Zielinski, Justin D. Shin, Shantanu P. Jadhav

## Abstract

Interactions between the hippocampus (area CA1) and prefrontal cortex (PFC) are crucial for memory-guided behavior. Theta oscillations (~8 Hz) underlie a key physiological mechanism for mediating these coordinated interactions, and theta oscillatory coherence and phase-locked spiking in the two regions have been shown to be important for spatial memory. Hippocampal place cell activity associated with theta oscillations encodes spatial position during behavior, and theta-phase associated spiking is known to further mediate a temporal code for space within CA1 place fields. Although prefrontal neurons are prominently phase-locked to hippocampal theta oscillations in spatial memory tasks, whether and how theta oscillations mediate processing of spatial information across these networks remains unclear. Here, we addressed these questions using simultaneous recordings of dorsal CA1 – PFC ensembles and population decoding analyses in male rats performing a continuous spatial working memory task known to require hippocampal-prefrontal interactions. We found that in addition to CA1, population activity in PFC can also encode the animal’s current spatial position on a theta-cycle timescale during memory-guided behavior. Coding of spatial position was coherent for CA1 and PFC ensembles, exhibiting correlated position representations within theta cycles. In addition, incorporating theta-phase information during decoding to account for theta-phase associated spiking resulted in a significant improvement in the accuracy of prefrontal spatial representations, similar to concurrent CA1 representations. These findings indicate a theta-oscillation mediated mechanism of temporal coordination for shared processing and communication of spatial information across the two networks during spatial memory-guided behavior.

## SIGNIFICANCE STATEMENT

Theta oscillation (~8 Hz) mediated interactions between the hippocampus and prefrontal cortex are known to be important for spatial memory. Hippocampal place-cell activity associated with theta oscillations underlies a rate and temporal code for spatial position, but it is not known whether these oscillations mediate simultaneous coding of spatial information in hippocampal-prefrontal networks. Here, we found that population activity in prefrontal cortex encodes animals’ current position coherently with hippocampal populations on a theta-cycle timescale. Further we found that theta-phase associated spiking significantly improves prefrontal coding of spatial position, in parallel with hippocampal coding. Our findings establish that theta oscillations mediate a temporal coordination mechanism for coherent coding of spatial position in hippocampal-prefrontal networks during memory-guided behavior.

## INTRODUCTION

The hippocampus and prefrontal cortex (PFC) are both necessary for learning and memory, and their interactions are critical for memory-guided behavior (Preston and Eichenbaum, 2013; Shin and Jadhav, 2016; Eichenbaum, 2017). Multiple inactivation studies in rodent spatial behavioral tasks have established a necessary role for functional interactions between the two regions (Floresco et al., 1997; Riedel et al., 1999; Wang and Cai, 2008; Churchwell et al., 2010; Maharjan et al., 2018). The physiological mechanisms that underlie these interactions, and how they support spatial memory processing, are thus key questions in the field.

Theta oscillations are prominent hippocampal local field potential oscillations (~8 Hz) associated with place-cell activity during spatial exploration in rodents, and have been established as a key physiological mechanism for mediating hippocampal (area CA1)-PFC interactions (Hyman et al., 2005; Jones and Wilson, 2005a; Siapas et al., 2005; Gordon, 2011; Shin and Jadhav, 2016). Theta-mediated CA1-PFC interactions play a crucial role in spatial memory tasks, and manifest as oscillatory coherence between theta rhythms in the two regions, as well as phase-locked spiking of neurons to hippocampal theta oscillations (Jones and Wilson, 2005a; Benchenane et al., 2010). Importantly, both oscillatory coherence and prefrontal phase-locked spiking have been shown to support performance in spatial memory tasks (Jones and Wilson, 2005a; Benchenane et al., 2010; Hyman et al., 2010; Hallock et al., 2016).

Theta-oscillation mediated hippocampal-prefrontal synchrony is therefore important for memory, but how this synchrony supports shared information processing to enable spatial memory behavior is still under investigation. There is evidence to suggest these interactions play a role in maintaining task-related representations for neurons in both regions (Jones and Wilson, 2005a; Hyman et al., 2011; Spellman et al., 2015), but several important questions about theta-mediated spatial representations remain unresolved. First, although spatial position is the most fundamental feature represented by hippocampal place-cell activity during theta oscillations (O’Keefe and Recce, 1993), whether spatial information is coherently encoded in the hippocampal-prefrontal network is not known. Spatial position during behavior can be accurately decoded from hippocampal ensemble activity (Wilson and McNaughton, 1993; Brown et al., 1998; Jensen and Lisman, 2000), and prefrontal neurons are known to have spatially selective representations (Jadhav et al., 2016; Mashhoori et al., 2018). However, no study has used simultaneous recordings and ensemble decoding to investigate whether hippocampal and prefrontal populations can coherently encode spatial position, given the evidence for strong theta-mediated interactions. Second, the role of theta phase-associated PFC spiking in spatial coding is not clear. It is known that in addition to a place-cell firing rate code in CA1, theta oscillations further underlie a temporal code for space based on theta phase precession, with the timing of a CA1 spike relative to the ongoing theta phase conveying finer information about where an animal is within a place field (O’Keefe and Recce, 1993; Skaggs et al., 1996; Harris et al., 2002; Schmidt et al., 2009). This phase-based temporal code significantly improves the encoding of spatial position by CA1 ensembles when theta phase is taken into account (Jensen and Lisman, 2000). Theta phase-locking as well as theta phase precession has been reported in prefrontal neurons (Jones and Wilson, 2005a, b; Hyman et al., 2010), but whether this phase-associated spiking leads to a similar improvement in spatial coding by prefrontal ensembles is not known.

We therefore addressed these questions using simultaneous hippocampal-prefrontal ensemble recordings and decoding analyses in rats performing a spatial memory task that requires their interactions (Maharjan et al., 2018). We find that prefrontal population activity can indeed encode the animal’s current position on a theta-cycle timescale coherently with CA1 coding, with correlated representations of position in the two regions. Further, we find that similar to CA1 representations, incorporation of theta phase also significantly improved prefrontal coding of spatial position, while maintaining coherent CA1-PFC coding. Our findings thus establish that theta oscillations mediate temporal coordination of hippocampal-prefrontal activity for coherent coding of spatial position during memory-guided behavior.

## MATERIALS AND METHODS

### Animals and experimental design

Five adult male Long Evans rats (450-550 g, 4-6 months, RRID: RGD_2308852) were used in this study. All procedures were conducted in accordance with the guidelines of the US National Institutes of Health and approved by the Institutional Animal Care and Use Committee at Brandeis University. Animals were individually housed and kept on a standard 12h/12h light-dark cycle, with initial *ad libitum* food and water prior to training and experimental protocols. Experimental protocols were carried out during animals’ light cycles. As previously described (Jadhav et al., 2016; Tang et al., 2017), after daily habituation and handling, animals were food deprived to 85-90% of their *ad libitum* weight and trained to seek liquid food reward (condensed milk) alternating between ends on an elevated linear track (80 cm, 7 cm wide track sections). Animals were also exposed and habituated to an elevated rest box during this training period. After animals reached a criterion level (50 rewards in 15-20 minute linear track sessions), they were taken off food restriction and subsequently, chronically implanted with a multi-tetrode drive (see *Surgical procedures, euthanasia, and histology*). Following recovery, animals were again food deprived to 85-90% of their *ad libitum* weight and re-habituated to the training task described above. Animals were then exposed to the full W-track continuous alternation behavioral task (see *Behavioral task*), and electrodes were positioned appropriately.

### Behavioral task

Rats performed a continuous W-track spatial alternation task that requires hippocampal-prefrontal interactions, as previously described (Jadhav et al., 2012; Jadhav et al., 2016; Tang et al., 2017; Maharjan et al., 2018). An experimental day consisted of multiple interleaved behavioral sessions on the W-track and rest box, consisting of 15-20 min sessions and 30-40 min rest periods respectively. The W-track (~80 x 80 cm) consists of elevated track sections (7 cm wide) with reward wells situated at the end of each of the three arms. The three arms were connected with two short sections (~40 cm long; Fig. 1). Calibrated evaporated milk rewards were delivered automatically via infrared detectors integrated in reward wells on correct trials. Rats were tasked with learning a continuous spatial alternation strategy (starting from the center arm), alternating visits to either side well (outbound component) and the center well (inbound component) (Figs. 1, 2). Incorrect alternations (visiting the same side well in consecutive outbound components – outbound error), incorrect side-to-side well visits (without visiting the center arm – inbound error), or perseverations (repeated visits to the same well just visited) were not rewarded. For each animal, the final two run sessions on the experimental day corresponding to high memory performance (87.0% ± 2.4% for outbound task phase; 94.0% ± 2.1% for inbound task phase) were used for analysis.

**Figure 1.**
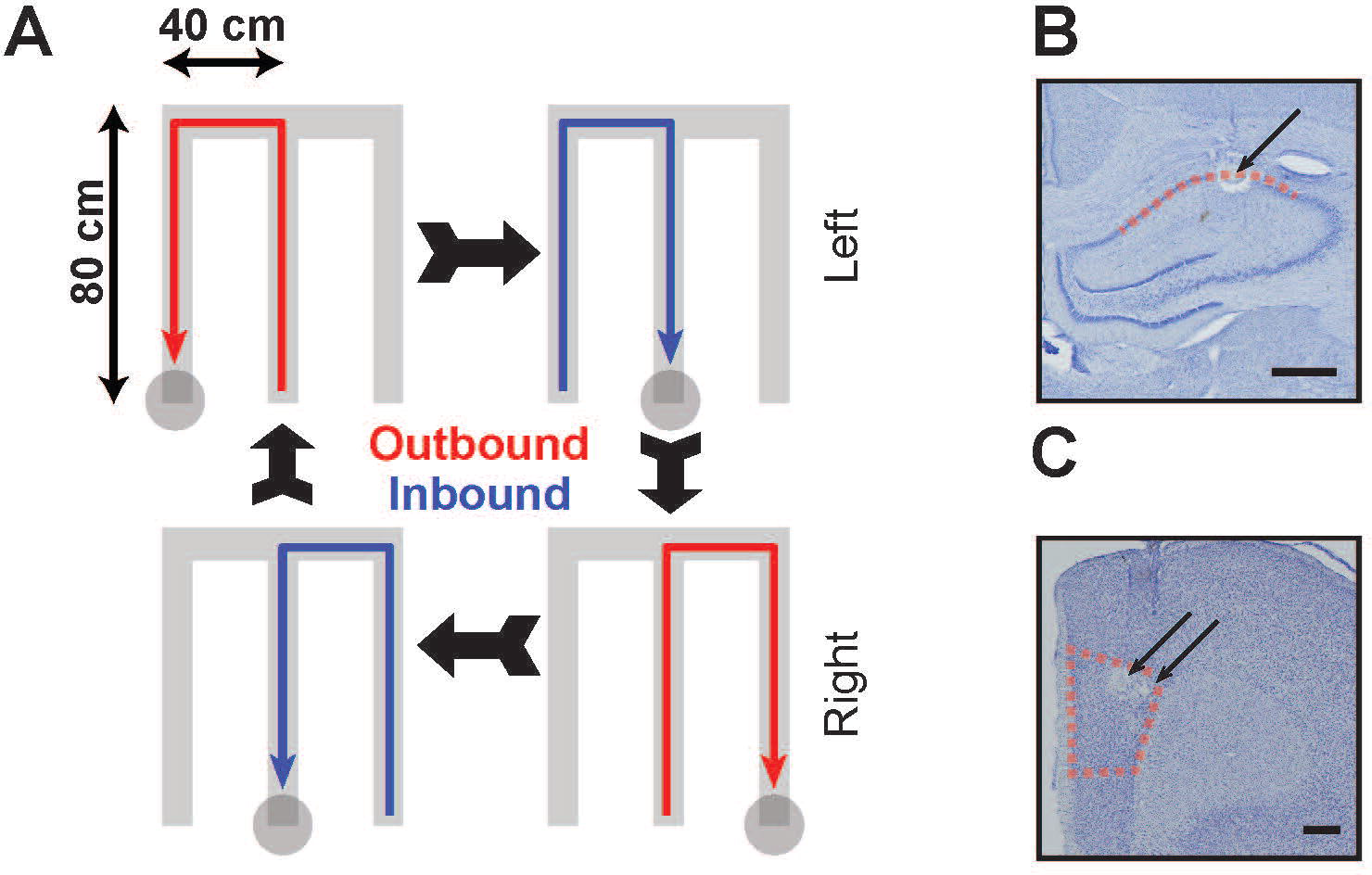
Experimental design and recording locations. **(A)** Schematic of W-track spatial alternation task. Animals run in a W-shaped maze, alternating between runs to the center arm and to the two alternating outer arms for liquid food reward (see **Materials and Methods** for more details). **(B-C)** Representative histological images of recording sites marked by lesions (arrows) in (**B**) dorsal CA1 and (**C**) medial PFC (prelimbic, or PrL, region). Boundaries indicate extent of tetrode locations identified in dorsal CA1 and PFC in all 5 animals. Scale bars indicate 500μm.

**Figure 2.**
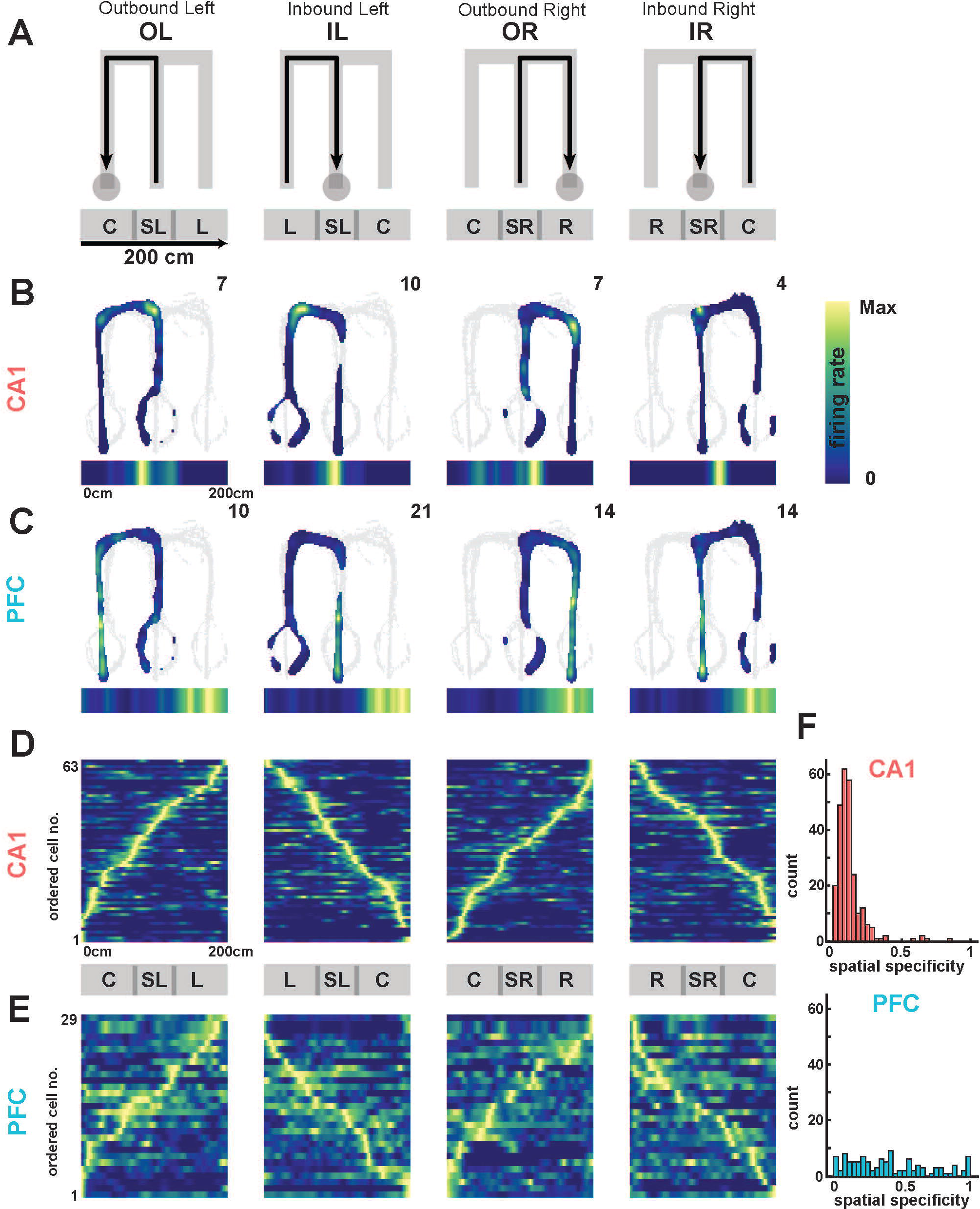
Spatial representation properties in CA1 and PFC. **(A)** Schematic of 2D firing rate maps as a function of W-track behavioral trajectories (From left to right: Outbound Left, or OL; Inbound Left, or IL; Outbound Right, or OR; Inbound Right, or IR). Below are schematic maps of linearized (1D) trajectories showing equivalent position with respect to the 2D behavior. Track arms are labeled as follows (C = Center, L = Left, SL = Side Left, R = Right, SR = Side Right). **(B)** Occupancy normalized spatial firing rate maps of an example CA1 unit for the four separate trajectory types. Peak firing rate is reported in the upper right, and linearized firing rate maps displayed below of each panel. **(C)** Same as in **(B)**, but for a representative PFC neuron. **(D)** Population of CA1 occupancy normalized linear firing rate maps for one animal, normalized and ordered by peak firing rate for each trajectory. Population activity mapped the spatial extent in all trajectories. **(E)** Same as in **(D)**, but for a representative PFC population in one animal. **(F)** Spatial specificity distributions in CA1 (*top*) and PFC (*bottom*) populations across all animals. Spatial specificity of CA1 neurons was significantly higher than PFC neurons (median spatial specificity (Inter-Quantile-Range, IQR); CA1, 0.10 (0.06 to 0.13); PFC, 0.37 (0.17 to 0.62); CA1 vs. PFC, Kruskal-Wallis test, p = 1e-99) Detailed statistics are reported in text.

### Surgical procedures, euthanasia, and histology

Surgical implantation procedures were as previously described (Jadhav et al., 2012; Jadhav et al., 2016; Tang et al., 2017), and post-operative analgesia administered for several days post implantation. Briefly, each rat was surgically implanted with a 3D printed microdrive containing 30 independently moveable tetrodes, with 15 tetrodes targeting right dorsal hippocampus (−3.6 mm AP and 2.2 mm ML) and 15 tetrodes targeting right PFC (+3.0 mm AP and 0.7mm ML). Tetrodes were made by twisting and bundling 4 NiCr wires (diameter 13 μm; Sandvik Palm Coast, Palm Coast, FL), followed by gold electroplating to an impedance of 200-300 kOhm. Electrodes were gradually advanced for 2-3 weeks following surgery to desired depths, concurrent with recovery and pre-training. The hippocampal layer was identified by characteristic LFP patterns such as presence of sharp-wave ripples (SWRs), SWR polarity and theta modulation.

At the end of the experiment, 24 hours prior to euthanasia, animals were anesthetized (1-2% isoflurane) and a current (30 μA) was passed through each tetrode to form lesions at their tips for localization. Animals were later euthanized (Beuthanasia 200 mg/kg) and perfused transcardially with 4% formaldehyde using approved procedures. Brains were then fixed in 4% formaldehyde and 30% sucrose, cut into 50-μm sections, stained with cresyl violet, and imaged for verification of tetrode localization.

### Electrophysiology and data acquisition

All tetrodes were referenced with respect to a cerebellar ground screw. For each animal, one tetrode in corpus callosum served as hippocampal reference electrode, and another tetrode in overlying cortical regions with no spiking signals served as prefrontal reference electrode. All behavioral and electrophysiological data was acquired using a SpikeGadgets system (SpikeGadgets, San Francisco, CA). Digital electrophysiological data was acquired using 128-channel digitizing headstages, sampled at 30 kHz and saved to disk, with spike data bandpass filtered between 600 Hz and 6 kHz, and local field potential (LFP) bandpass filtered between 0.5 Hz to 400 Hz and down sampled to 1.5 kHz. Input and output triggers for behavioral and environmental data (eg. reward delivery) were recorded at 1-ms resolution and synchronized to electrophysiological data. Animal movement and behavior was recorded and tracked using an overhead color CCD camera (30 fps), with animal head position indicated by color LEDs affixed to the headstage apparatus and microdrive. Cameras were calibrated to provide a resolution of 0.1 cm/pixel, and spatial extent of LEDs permitted a tracking resolution of ~2 cm.

### Unit identification and inclusion

Spike peaks were identified by a threshold crossing of 40 μV in the filtered spike band for hippocampus (CA1) and prefrontal cortex (PFC) respectively. Spikes were manually sorted as previously described (Jadhav et al., 2012; Jadhav et al., 2016; Tang et al., 2017). Briefly, putative spikes had clustering parameters extracted (spike width on each channel, spike amplitude, and principal components), and were clustered using a custom Matlab (MathWorks, Natick, MA; RRID: SCR_001622) cluster visualization program (MatClust). Clusters were judged based on waveform shape, isolation distance, and lack of ISI violation. Only well isolated and stable putative excitatory units across the sessions were included, with putative interneurons identified and excluded based on average firing rate ≥ 15 Hz and spike width criteria, as previously described (Jadhav et al., 2016; Tang et al., 2017). Further, only neurons which fired at least 100 spikes in each session were included for further analysis.

### Spatial maps and linearization

Two-dimensional occupancy-normalized spatial firing maps were calculated for each unit when the animals’ speed was greater than 3 cm/s, with spikes binned in 1 cm square bins and smoothed with a 2D Gaussian (2 σ), excluding spiking during high ripple power (>3 SD ripple band power, see *LFP collection and high-theta segmentation*). The linearized spiking activity of each cell was computed by assigning the rat’s linear position along the 2D skeleton of the four possible linear behavioral trajectories (center arm reward well to outer arm reward well for outbound trajectories, and the converse for inbound trajectories; (Frank et al., 2000; Jadhav et al., 2016)). Spiking and occupancy closest to each linear 2 cm bin on these four trajectories was then used to calculate the smoothed, occupancy-normalized linear firing rate for correct trajectories. A peak rate of 3 Hz or greater was required for a cell to be considered a place cell in CA1, and a similar criterion was applied to PFC cells for inclusion in analysis (Jadhav et al., 2016). These linearized trajectories with occupancy-normalized firing rates were used in all subsequent decoding analyses.

### Spatial specificity and spatial coverage

Concatenating the 1D firing rate maps for all four behavioral trajectories, spatial specificity was computed as the spatial sparsity of firing (Fig. 2), the proportion of firing rate bins greater than 25% of the peak firing rate (Jadhav et al., 2016). Spatial specificities varied from 0 for highly spatially specific neurons, to 1 for neurons with uniform firing fields (range of spatial specificity: 0-1). Spatial coverage of a trajectory by an ensemble was calculated as the percentage of spatial bins with population occupancy normalized firing rate ≥ 3 Hz (Kay et al., 2016).

### LFP collection and high-theta segmentation

LFP was band pass filtered in the delta (1-4 Hz), theta (6-12 Hz), and ripple bands (150-250 Hz) using zero phase IIR Butterworth filters. We determined envelopes and phases by Hilbert transform, and took the ratio of the theta to delta envelopes at each time point for every hippocampal tetrode. High theta periods were detected using criteria for theta power, running speed, and exclusion of sharp-wave ripples (SWRs). Specifically, high theta windows were assigned as time periods at least one theta cycle long when the smoothed (1 σ) mean theta/delta ratio exceeded 2, no SWRs were detected with a 3 SD threshold in the absolute power of the ripple band, and animal speed was greater than 3 cm/s.

### Theta Coherence

Coherence was calculated for each trajectory type using the Chronux (http://chronux.org/, RRID:SCR_005547) spectral processing package for MATLAB as follows. Saturating movement artifacts in the LFP were removed, and for valid time periods in which the animal’s speed was greater than 3 cm/s, the animal’s linear position was interpolated to the nearest 4 cm linear bin. The LFP of the PFC and CA1 tetrodes with the highest number of neurons in a given session were used to calculate coherence between the two regions. Each coherogram time bin was then assigned to a particular linear bin with the closest spatial position based on the animal’s movement, leading to the average coherogram across trajectories as a function of linear distance (Fig. 3). All further analyses were averaged across animal and session for display.

**Figure 3.**
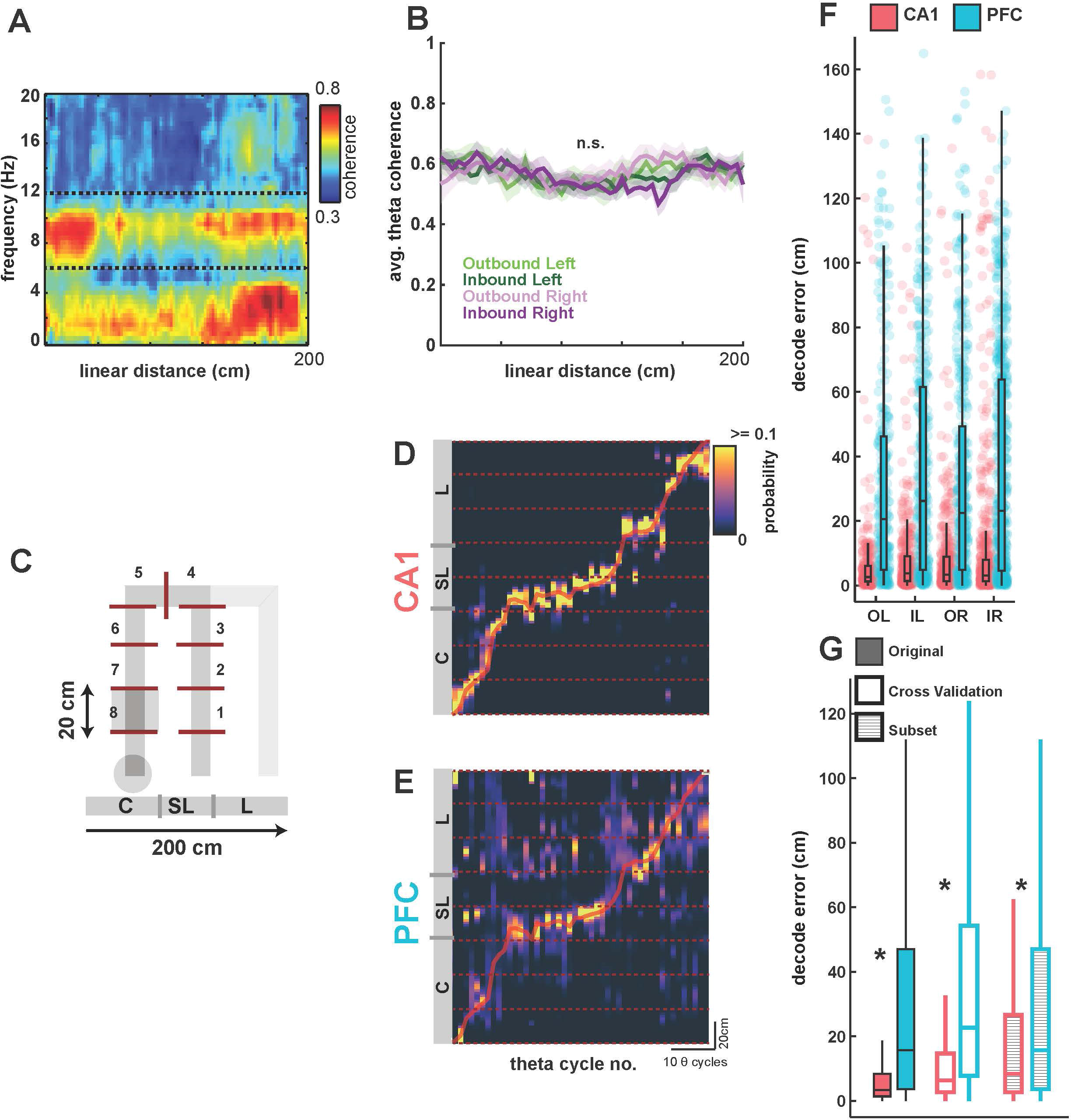
Decoding of spatial position from CA1 and PFC ensembles on a theta-cycle timescale. **(A)** Average cohereogram plotted against linear position between CA1 and PFC tetrodes for all Outbound Left (OL) trajectories across animals. Note the high coherence in the theta range throughout the memory-guided trajectory. **(B)** Average coherence in the theta band for all animals, for the four trajectory types, with s.e.m. overlaid. Theta coherence was equivalently high for all trajectory types (p > 0.63). **(C)** Schematic of spatial binning of trajectories, illustrated for Outbound Left. Each trajectory was split into 20 cm bins (or, 8 spatial sections) for further analysis, excluding the sections with reward wells. All four trajectory types were similarly divided into 8 sections. **(D)** Actual spatial position of animal (red line) during an outbound left trajectory is shown along-with Bayesian decoded spatial position (heatmap) from CA1 ensembles (n = 63 neurons). Decoded position is computed as peak probability of decoded position in theta cycle time bins, using underlying theta cycles during the course of the trajectory (median length of theta cycles is 131 ms). Theta cycle number is indicated along the X-axis, and position along the Y-axis is shown across the eight 20-cm spatial sections illustrated in **C**. **(E)** Same as in **(D)**, but showing simultaneous Bayesian decoding of position using PFC ensembles (n = 29 neurons). Heatmap is similar to **D**. **(F)** Distributions of decoding error (difference between actual and decoded position) in theta-cycle time bins (actual values as dots, median and inter-quartile ranges overlaid) are shown for both CA1 and PFC ensemble decoding for all four trajectory types for one session from one animal (*N* = 282, 357, 417 and 569 time bins respectively for the four trajectory types; OL, IL, OR and IR). CA1 ensembles had significantly better decoding accuracy than PFC ensembles for all trajectory types (all p ≤ 1e-33; detailed statistics in text). **(G)** Comparison of CA1 and PFC decoding error under different decoding conditions, averaged across all trajectory types for all sessions. Filled bars as in **F**; open bars using leave-one-trial-out cross-validation (LOOCV) for decoding, hatched bars using sub-sampling to match the number of CA1 and PFC neurons for decoding. CA1 had significantly smaller decoding error under all conditions (* = p < 3e-60; detailed statistics in text).

### Theta cycles theta phase, and decoding

Theta LFP phase for both hippocampal and prefrontal unit spiking was relative to a hippocampal reference tetrode located in corpus callosum, as previously described (Jadhav et al., 2016). Within high theta time windows detected as described above, the troughs of the hippocampal theta-filtered LFP were identified and used to segment valid theta cycles, discarding cycles where phase was ambiguous or reset, as previously described (Johnson and Redish; Gupta et al., 2012; Feng et al., 2015; Wu et al., 2017). Only theta cycles with at least 3 simultaneously active template cells in both hippocampus and prefrontal cortex were analyzed, and theta cycle decoding was implemented as previously described (Johnson and Redish; Gupta et al., 2012; Wu et al., 2017). A Bayesian decoder was used to calculate the probability of the animal’s location given the firing rate templates of the neurons that fired and their spikes that occurred in each time window (Zhang et al., 1998; Davidson et al., 2009; Karlsson and Frank, 2009; Gupta et al., 2012; Pfeiffer and Foster, 2013). Briefly, the probability of the animal’s position (*pos*) across all total spatial bins (*S*) given a time window (*t*), containing Poisson spiking (*spikes*) of independent units is

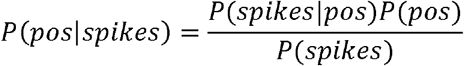

Normalizing over *P*(*spikes*) and using a uniform prior *P*(*pos*) to avoid spatial decoding bias, we get

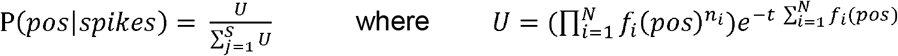

Where *f*_*i*_(*pos*) is the occupancy normalized 1D firing rate map for the *i*-th unit, *N* is the total number of active units, *n*_*i*_ is the number of spikes fired by a particular *i*-th unit, and *t* is a time window (in this case, the entire theta cycle). This was computed using the firing rate template for the corresponding behavioral trajectories. The decoded position is then the non-zero spatial bin location with the maximum posterior probability. Decoding error is defined as the absolute difference between the animal’s actual position averaged in the time window and the decoded position (Fig. 3).

### Cross validation metrics – firing rate templates and neuron count

Cross validation procedures were used as described (Fig. 3G) to generate null distributions for significance testing. To verify that the templates constructed across trials were sufficiently powered for subsequent population decoding, we used a standard leave one (trial) out cross validation strategy (LOOCV) (van der Meer et al., 2017). Briefly, the firing of both populations during one randomly selected trajectory were left out during field/template construction, with template construction then proceeding as described above with the remaining trials. The animal position for theta cycles during this omitted trial was then decoded using these cross-validation templates, and we repeated this LOOCV procedure for all trials per session.

Next, to confirm that decoding accuracy in CA1 was not simply due to a higher isolated neuron count (Fig. 3G), we confirmed our results by randomly sub-setting our CA1 neuron count to match the PFC neuron count in each session and animal (number of CA1 cells was always higher than PFC cells, Table 1) and proceeding with the spatial decoding procedure as above. We repeated this procedure 1000 times per session, each with a random choice for CA1 neural subsets. For the random CA1 neuron subsets, we preserved spatial coverage and spatial specificity in CA1 (to avoid artifacts in spatial decoding due to incomplete coverage) by drawing from the CA1 neuron distribution weighted by the 20-cm spatial section in which their peak firing rate occurred.

### Joint decoding in CA1 and PFC

Using previously described methods (Haggerty and Ji, 2015; Saleem et al., 2018), joint decoding error in each session (*n* = 10 sessions in 5 animals) was quantified as the Pearson’s linear correlation coefficient between the two decoded position errors in CA1 and PFC within each time window, computed for each 20-cm spatial section/bin. Animal speed at each time window was classified into three evenly spaced speed bins: low, medium, and high; 3-15, 15-30, and 30+ cm/s respectively (we obtained similar results with other thresholds for speed categories, including 3-12, 12-24, 24+ cm/s, and 3-17, 17-34, 34+ cm/s), and correlation was computed for decoding errors within each 20-cm spatial section/bin (20-180 cm in 20 cm increments for each spatial bin), for subsequent comparison to shuffled controls while matching speed category and position bin. Significance for joint decoding error was computed relative to shuffled speed and position bin matched time windows, repeating the above procedure 1000 times, ensuring that correlations in decode error due to speed and location were controlled for. Joint decode errors for both real and shuffled data were binned in 1cm^2^ bins and smoothed with a 2σ 20 cm gaussian and z-scored for visualization, with the residual joint decoding computed as the difference between the two (actual – shuffle; Fig. 4).

**Figure 4.**
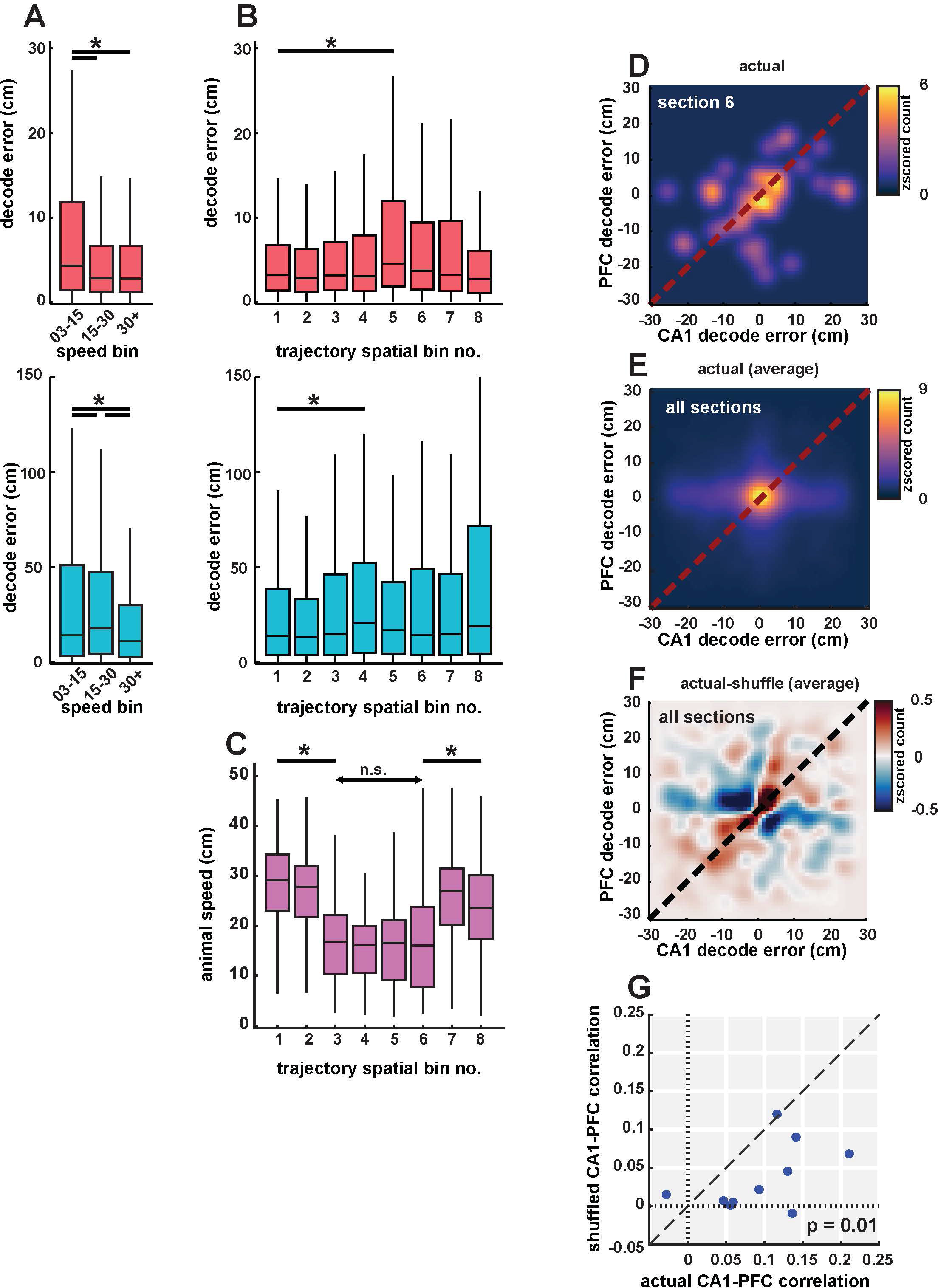
Coherent spatial position coding in CA1 and PFC. **(A)** Decoding error from CA1 (pink, *top*) and PFC (cyan, *bottom*) populations in low (3-15 cm/sec), medium (15-30 cm/sec) and high speed (30+ cm/sec) categories. Decoding error varied with speed in both areas. **(B)** Decoding error as a function of spatial bin (sections) for CA1 (pink, *top*) and PFC (cyan, *bottom*). Decoding error had a small but significant increase in middle bins (bin 4 and 5). **(C)** Animal speed as a function of spatial bin. Animal speed decreased significantly in middle of trajectories corresponding to turn and choice point sections. **(D)** 2-d histogram showing cycle-by-cycle correlations between CA1 and PFC decoding errors in spatial section 6. Heatmap indicates z-scored histogram count of decoding error for CA1 vs. PFC. **(E)** Same as **D**, averaged across all spatial bins. Decoding error distribution showed peaks along the diagonal around 0, with significant correlations for CA1 vs. PFC decoding error (r = 0.10, n = 10 sessions, p = 0.001). **(F)** Difference between decoding errors using actual and shuffled data (controlled for spatial and speed bins) shows a distribution along the diagonal, indicating strong correlations between actual decoded errors. **(G)** Decoding error correlations for actual vs. shuffled data for the 10 recording sessions. Actual correlations were significantly higher than shuffled values (mean correlation values of r = 0.10 vs. r = 0.03, p = 0.01). Detailed statistics for all comparisons are reported in text.

### Phase locking, concentration, and precession

Phase locking (Fig. 5) was computed using a Rayleigh z-test for non-uniformity using the spikes of all neurons for both sessions per animal. As previously described (Jadhav et al., 2016), time periods with animal speed > 3 cm/s and with absolute power in the ripple band was less than 3 SD were considered, with spike phase derived from the hippocampal reference tetrode. Neurons with a criterion significance level of p < 0.05 were considered significantly phase locked. Concentration parameters (kappa) were computed for phase locked neurons via a von-Mises fit to the spiking phase data as subset above. Phase precession was computed using same criterion for neurons and spikes as above, with the addition of limiting the phase precession analysis to the peak firing field, computed as the linear distance from the peak firing rate to the first instance of firing rate < 0.25 of the peak firing rate in both directions. Phase precession values were then computed as previously described (Kempter et al., 2012; Cei et al., 2014; Jeewajee et al., 2014) (See https://github.com/HoniSanders/measure_phaseprec for publicly available code). Briefly, we computed the best fit phase slope and phase intercept of a linear-circular distribution over spiking linear distance and spiking phase, respectively. Slope was limited to a maximum of one cycle over the peak firing field to reduce overfitting.

**Figure 5.**
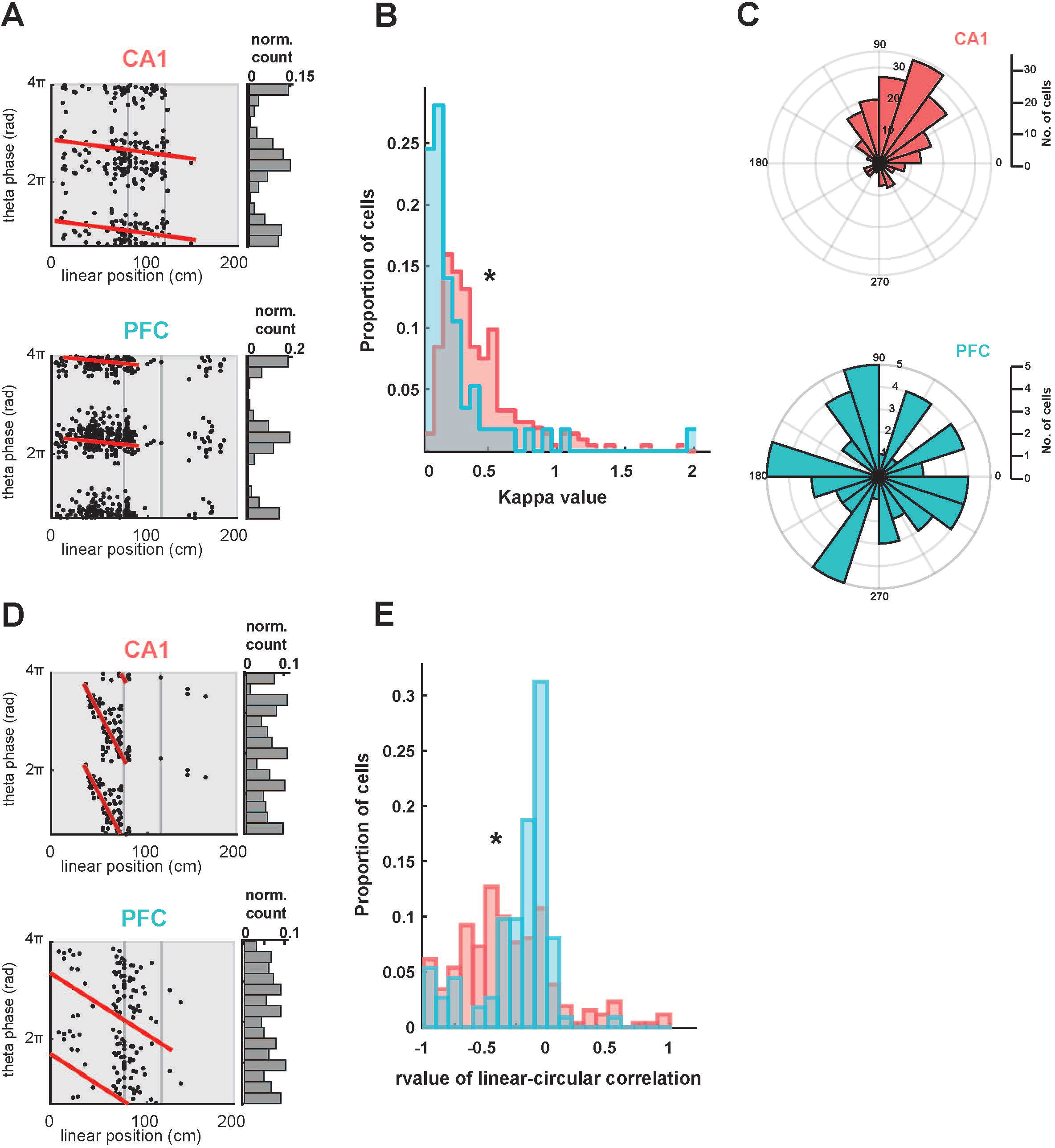
Theta phase modulation of CA1 and PFC spiking. **(A)** Examples of theta phase-locked CA1 and PFC neurons. Spikes from single neurons are plotted with corresponding theta oscillation phase and linear position indicated along the two axes. Note that spikes occur at specific phases of theta oscillations within the firing fields of the neurons, thus exhibiting a phase-space relationship. Both neurons were significantly phase-locked, assessed with a Rayleigh z-test (CA1 neuron, kappa = 1.1 p = 1.6e-28; PFC neuron, kappa = 2.0, p = 1e-99). These neurons did not show significant phase precession (p > 0.63, red lines). **(B)** Distribution of kappa values for all phase-locked CA1 (n = 213) and PFC (n = 57) neurons. CA1 had significantly stronger phase-locking than PFC (p = 1.5e-08). **(C)** Distribution of preferred theta phases for significantly phase locked CA1 (n = 213) and PFC (n = 57) neurons. The two distributions are significantly different from each other (p = 0.002). **(D)** Examples of theta phase-precession in CA1 and PFC neurons. Spikes from single neurons are plotted with corresponding theta oscillation phase and linear position indicated along the two axes, with line fits for circular-linear correlation overlaid in red (CA1 neuron, ρ = −0.64, p = 1.7e-7; PFC neuron, ρ = −0.27, p = 0.02). **(E)** Distribution of rho-value (ρ) from line fits for circular-linear correlation quantifying strength of phase precession in all CA1 and PFC neurons. CA1 had significantly stronger phase precession than PFC (p = 2.7e-4).

### Theta phase splits and shuffling

For analyses involving multiple phase bins (Fig. 6), the following procedure was followed. Each theta cycle used for analysis was divided into *N* evenly spaced sub-periods divided by phase of the theta period, resulting in *N* different spiking templates defined by theta phase for further Bayesian reconstruction. Templates were then used for every cell that corresponded to its instantaneous spike phase in theta. Shuffling protocol consisted of randomly assigning spike phase bins during template construction and decoding normally as described above.

**Figure 6.**
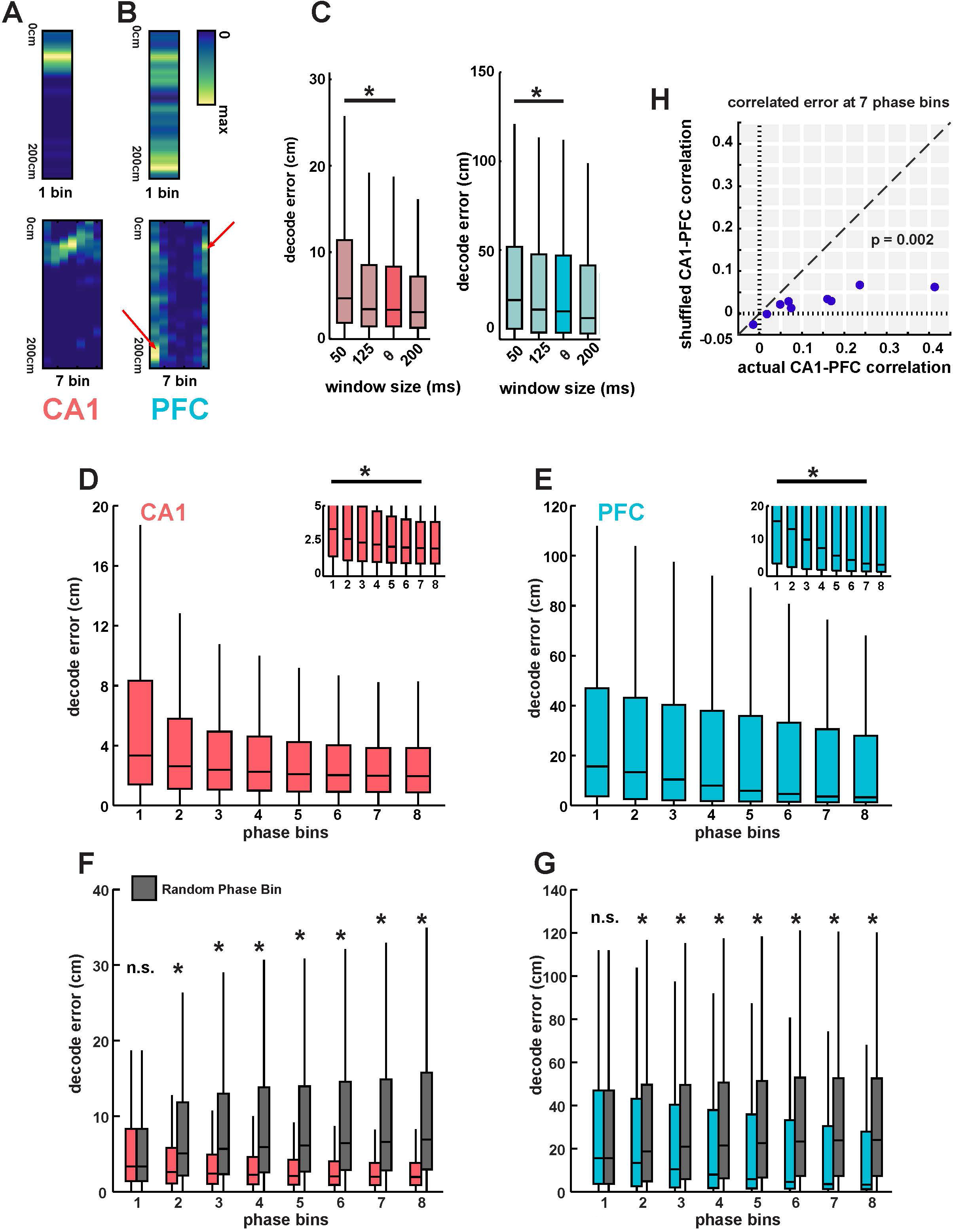
Theta phase-based templates improves spatial position decoding in both CA1 and PFC. **(A)** Example of a CA1 firing field split into 7 theta phase-based templates. This neuron exhibited strong phase precession, apparent in the phase-based templates shown on right, where preferred firing location changes with theta phase. **(B)** Example of a PFC firing field split into 7 theta phase-based templates. This neuron exhibited strong phase locking, apparent in the phase-based templates shown on right, where only preferred phase bins (indicated by arrows) shows significant spiking that is spatially localized. Non-preferred phase bins exhibit highly reduced number spikes. **(C)** Decoding error using firing rate templates as a function of time windows. Using smaller temporal windows (50 ms) than theta cycles leads to increased decoding error (p < 3.7e-9). **(D)** Decoding error using CA1 ensembles using successively increasing number of phase-bin templates. Phase bin 1 corresponds to a firing-rate-only template, similar to Fig. 3. Decoding accuracy significantly increased until 5 phase bins (p ≤ 0.02, ~2-fold improvement in median decoding accuracy), with only non-significant, asymptomatic improvement with subsequent increase in phase bins. Inset shows low decoding error values for clarity. **(E)** Same as **D**, using PFC ensembles. Decoding accuracy significantly increased until 7 phase bins (p ≤ 0.0013, ~4-fold improvement in median decoding accuracy). **(F)** Comparison of decoding error using phase-based templates to an equivalent number of templates, but with random assignment of theta phase (random phase bin). Decoding error with shuffled templates using random phase assignment did not show any significant improvement in decoding accuracy with increase in phase bins (p < 3e-34 for all pairwise comparisons from 2-8 phase bins between actual and shuffled decoding values). **(G)** Decoding error correlations for actual vs. shuffled data for the 10 recording sessions for 7 phase bins. Actual correlations were significantly higher than shuffled values (mean correlation values of r = 0.13 vs. r = 0.02, p = 0.009).

## RESULTS

We used multi-site multi-electrode recordings to simultaneously record neural activity in dorsal CA1 region of hippocampus and medial prefrontal cortex (PFC) of rats (n = 5) as they performed a continuous W-track spatial alternation task that requires hippocampal-prefrontal interactions (Kim and Frank, 2009; Jadhav et al., 2012; Jadhav et al., 2016; Tang et al., 2017; Maharjan et al., 2018). Fig. 1 illustrates a schematic of the task and histologically verified recording locations in CA1 and PFC. Recordings in all animals were localized primarily to dorsal CA1 region and the PreLimbic (PrL) area of PFC, delineated in Fig. 1B, C. The W-track alternation task (Fig. 1A) requires animals to visit the two outer arms of the maze in an alternating sequence, with rewarded visits from the center arm to alternating outer arms (2 outbound trajectories) interleaved with rewarded returns from the outer arms to the starting center arm (2 inbound trajectories) (see **Materials and Methods**). Data from two behavior sessions per animal were used for analyses, with animals performing the task with high accuracy in each session (n = 10 sessions; n = 32.6 ± 2.8 correct trials per session, performance: outbound task phase, mean ±□s.e.m; 87.0% ± 2.4%; inbound task phase; 94.0% ± 2.1% inbound task phase). A total of 255 CA1 (animal mean ±□s.e.m; 51 ± 4) and 111 PFC (animal mean ±□s.e.m; 22 ± 3) neurons were used after exclusion of putative interneurons and those with less than 100 spikes (Table 1 shows distribution of neurons across 5 animals, with 2 sessions per animal with the same number of neurons in each session).

**Table 1.**

| HPC | PFC |
| --- | --- |
| 62 | 29 |
| 53 | 25 |
| 43 | 25 |
| 40 | 13 |
| 57 | 19 |

### Spatial representations in CA1 and PFC

Similar to CA1 place cells, PFC neurons are also reported to have spatially restricted, trajectory-specific firing in spatial choice tasks (Fujisawa et al., 2008; Ito et al., 2015; Jadhav et al., 2016; Mashhoori et al., 2018). Fig. 2A-C show illustrative CA1 and PFC neurons with spatially specific firing fields on the 4 trajectories of the W-maze task, namely, outbound to left arm (OL), inbound from left arm (IL), outbound from right arm (OR), inbound from right arm (IR). The four trajectories in the task are illustrated in Fig. 2A, with the maze divided into central arm (C), left arm (L), right arm (R), and two short sections connecting the arms, side-left (SL) and side-right (SR). Each trajectory consists of a run through from one reward well to another through 3 sections of the maze, e.g. outbound left (OL) through Center, Side-Left and Left Arm. Spatial firing fields for each CA1 and PFC neuron were computed as occupancy-normalized firing rates for each trajectory type, and then subsequently linearized for elapsed trajectory distance, similar to previous studies ((Frank et al., 2000; Jadhav et al., 2016); bottom panels in Fig. 2B, C show linearized fields with a 2-cm bin size; see **Materials and Methods** for details).

Since CA1 and PFC neurons have trajectory-specific firing fields (McNaughton et al., 1983; Frank et al., 2000; Hok et al., 2005; Fujisawa et al., 2008; Spellman et al., 2015), each trajectory is expected to have distinct ensemble representations or spatial templates. We therefore examined the response profiles of neurons in both regions across the four behavioral trajectories (example for one animal in Fig. 2D, E). Ensemble representations in both areas tiled the full extent of the spatial environment across all trajectories, with spatial coverage sufficient for subsequent decoding analyses for all animals, calculated as the percentage of spatial bins with population occupancy normalized firing rate ≥ 3 Hz ((Kay et al., 2016); mean % coverage ± SD; CA1, 97% ± 0.88%; PFC, 100% ± 0%). Note that neurons illustrated in Fig. 2D, E are sequentially ordered separately for each trajectory based on the position of their peak firing fields within that trajectory.

The spatial firing properties of CA1 and PFC were quantified by computing the spatial specificity (or, spatial sparsity) of neurons in both areas as previously described (Jadhav et al., 2016). Spatial specificity was defined as the fraction of each trajectory with above-threshold occupancy normalized firing rates (> 25% peak firing rate for each neuron, **Materials and Methods**). As expected, CA1 neurons had higher spatial specificity than PFC neurons, with the CA1 distribution skewed toward lower values of spatial specificity characteristic of highly place specific cell firing, and a spread of spatial specificities in PFC (Fig. 2E, Wilcoxon signed-rank test compared to uniform distributions with median = 0.5; CA1: Z = −13.8, p = 3.1e-43; PFC: Z = −3.0, p = 0.003). Spatial specificity of CA1 neurons was significantly higher than PFC neurons (median spatial specificity (Inter Quantile Range or IQR); CA1: 0.10 (0.06 to 0.13); PFC: 0.37 (0.17 to 0.62); CA1 vs. PFC spatial specificity distributions are significantly different from one another, Kruskal-Wallis test; Χ^2^(1) = 101.6, p = 1e-99), therefore with a ~4-fold ratio of median spatial specificities for CA1 vs. PFC neurons.

### CA1 and PFC populations encode spatial position at a theta cycle timescale

It is well-established that theta oscillations and theta coherence are prominent in hippocampal-prefrontal networks during memory-guided trajectories (Jones and Wilson, 2005a; Siapas et al., 2005; Benchenane et al., 2010; Hyman et al., 2010; Remondes and Wilson, 2013). We similarly observed high coherence between theta oscillations for all outbound and inbound trajectories. Fig. 3A illustrates a coherogram exhibiting high coherence in the theta band between CA1 and PFC electrodes for a representative outbound left trajectory, and Fig. 3B shows average theta coherence across sessions for the four trajectories. Theta coherence was similar across trajectories (average theta coherence for entire trajectory; Kruskal-Wallis with Bonferroni *post-hoc* for comparisons between the four trajectory types; Χ^2^(3) = 3.88, all comparisons p > 0.63).

It has been proposed that high theta coherence is suggestive of interactions on a theta oscillation timescale between hippocampal and prefrontal activity. We therefore asked, with what accuracy could we determine an animal’s current position during trajectories from the concurrent neuronal ensemble activity in either area, using theta oscillations as time bins? To address this question, we used a Bayesian decoder (see **Materials and Methods**) to estimate the probability of an animal’s position per time bin given the ensemble firing in the corresponding trajectory template, similar to previous studies (Brown et al., 1998; Jensen and Lisman, 2000).

We used the population templates in CA1 and PFC (Fig. 2D, E) corresponding to an animal’s current trajectory for all subsequent analyses, and corresponding theta oscillation cycles as time bins (see also **Materials and Methods**). Theta cycles have been proposed to represent temporal windows for organization of hippocampal-cortical activity (Itskov et al., 2008; Lisman and Redish, 2009), and previous studies have used theta cycle time bins for decoding CA1 ensemble activity (Gupta et al., 2012; Feng et al., 2015; Wang et al., 2015; Wikenheiser and Redish; Wu et al., 2017). We therefore used a similar approach for simultaneous decoding of CA1 and PFC activity. Decoded position was estimated for all four trajectories which were divided into eight 20-cm spatial bins or sections from the start to the end of the trajectory (Fig. 3C). We excluded the start and end sections of each arm with reward wells where animals are primarily stationary and hippocampal activity prominently exhibits sharp-wave ripples (Karlsson and Frank, 2009; Jadhav et al., 2016). Using this decoding analysis approach, we asked whether both CA1 and PFC ensembles represented current spatial position on a theta time-scale, and the degree of correlation, if any, in decoding errors (as a measure of position estimate accuracy) in the two areas.

Theta cycles were extracted from filtered LFP (6-12 Hz, see **Materials and Methods**), and segmented based on reference cycle troughs, similar to segmentation protocols in previous studies ((Johnson and Redish; Gupta et al., 2012; Feng et al., 2015; Wu et al., 2017); length of theta cycles: median (IQR), 131 ms (121.3 to 142.7 ms)). An example of decoded positions for an outbound left trajectory, relative to actual position, using CA1 and PFC ensembles is shown in Fig. 3D, E. The spatial bin with the highest non-zero posterior probability in a given time frame (theta cycle time) is assigned as the decoded position, and the animal’s mean linearized position during the same time window is the actual position of the animal. Decoding error is reported as the difference, in cm, between the actual and decoded position. Distribution of decoding error per theta cycle time bin using CA1 and PFC ensembles for all four trajectory types (outbound and inbound left (OL and IL, respectively), outbound and inbound right (OR and IR, respectively)) across all trials for one animal is shown in Fig. 3F (N = 282, 357, 417 and 569 time bins respectively in N = 10, 14, 12 and 10 trials in this session for the four trajectory types; OL, IL, OR and IR). For CA1 ensembles, the decoding error distribution is highly skewed to low values indicating high spatial decoding accuracy, as expected from previous studies (Jensen and Lisman, 2000) (range of medians and IQRs for CA1 decoding error across all animals and trajectories, 2.6-4.7 cm (4.4-11.5cm), Shapiro-Wilk test of normality showing non-normal distributions; p-values for all animals and trajectories ≤ 3.4e-10). We also observed the same skewed trend toward spatial decoding accuracy in PFC (range of medians and IQRs for PFC decoding error across all animals and trajectories, 5.1-28.9 cm (12.0-59.7 cm), Shapiro-Wilk test of normality showing non-normal distributions; p-values for all animals and trajectories ≤ 2.8e-08). Decoding error across the four trajectory types did not show differences for either CA1 or PFC populations (Kruskal-Wallis with Bonferroni correction for multiple comparisons; CA1, Χ^2^(3) = 7.04, all pairwise comparisons p ≥ 0.05; PFC, Χ^2^(3) = 2.55, all pairwise comparisons p ≥ 0.88), and importantly, CA1 decoding error was significantly smaller than PFC for all four trajectory types (Wilcoxon rank sum for all trajectories; Z ≤ −12.01, all comparisons p≤1e-33).

We next combined decoding errors across all trajectory types to compare and quantify CA1 and PFC decoding accuracy under different decoding conditions. Fig. 3G shows median decoding error using CA1 and PFC ensembles for all animals and trajectories (N = 93, 61, 52, 68, 52 correct trajectories for the 5 animals over both sessions). Both CA1 and PFC activity can be used to decode animal’s current position with relatively high decoding accuracy (solid bars, median (IQR); CA1: 3.4 cm (1.4 to 8.3 cm); PFC: 15.7 cm (3.7 to 47.0 cm)), with significantly more accurate estimation for CA1 (CA1 vs. PFC, Kruskal-Wallis, Χ^2^(1) = 2504.64, p ≤ 1e-99). Ratio of median decoding accuracy in the two areas was comparable to spatial specificity ratios shown in Fig. 1 (median decoding error ratio for CA1 vs. PFC ~4.6:1 for these decoding parameters; spatial specificity ratio for CA1 vs. PFC is ~4:1). We also used a “leave-one-trial-out” cross-validation approach for decoding (LOOCV), in which for decoding a given trajectory, templates used for decoding were built by omitting that trajectory ((van der Meer et al., 2017); see **Materials and Methods**), and observed similar results (open bars in Fig. 3G; median (IQR); CA1: 6.32 cm (2.7 to 14.7 cm); PFC: 22.7 cm (7.8 to 54.1 cm); CA1 vs. PFC, Kruskal-Wallis, Χ^2^(1)= 9333.2, p = 1e-99; median decoding error ratio for CA1 vs. PFC is ~4:1). The cross-validated decoding confirms that templates were sufficiently stable across trials to provide observed high decoding accuracy. Finally, in order to account for the difference in number of neurons for CA1 vs. PFC, we sub-sampled the number of CA1 neurons to match the number of PFC neurons for each session (hatched bars in Fig. 3G, note that PFC decoding errors are same as original data with solid bars). The median decoding error ratio improved since the reduction in number of CA1 neurons increased CA1 decoding error, but the CA1 ensembles still had significantly higher decoding accuracy than PFC ensembles (hatched bars in Fig. 3G; median (IQR); CA1: 8.4 cm (2.7 to 26.6 cm); PFC: 15.7 cm (3.7 to 47.0 cm); CA1 subset vs. PFC, Kruskal-Wallis, Χ^2^(1) = 272.97, p = 2.6e-61; median decoding error ratio for CA1 vs. PFC is ~2:1).

### Coherent representation of spatial position by CA1 and PFC ensembles

In order to examine whether representation of spatial position is shared across hippocampus and prefrontal cortex, we first examined the effect of behavioral variables, including running speed, on decoding accuracy. Running speed is an important variable that can vary from trial-to-trial and within trajectories, and since previous results have linked theta power and spatial representations with animal running speed (McFarland et al., 1975; McNaughton et al., 1983; Geisler et al., 2007; Maurer et al., 2012), we also analyzed decoding error with regard to animal speed within a given theta window. Each theta-cycle time bin was assigned low, medium and high speed ranges, similar to previous studies (Saleem et al., 2018) (3-15cm/sec, 15-30cm/sec, and >30 cm/sec respectively; we observed similar effect for other thresholds for speed categories with boundaries at 12 and 24cm/s, as well as 17 and 34 cm/s), and decoding error was compared for different speed categories. As expected, we saw a change in decoding error with speed, especially with an increase in CA1 decoding error for lower speeds (Fig. 4A, Kruskal-Wallis with Bonferroni *post-hoc*; CA1, Χ^2^(2) = 128.5, p ≤ 7.8e-15 for low vs. mid and low vs. high speed categories; PFC, Χ^2^(2) = 50.2, p ≤ 0.008 for low vs mid, low vs high, and mid vs. high speed categories). Examining the effect of spatial bins within trajectories, we also saw a small but significant increase in decoding error at the choice point and turn sections (spatial bins 4-5 on side-left and side-right arms, as illustrated in Fig. 3C) corresponding to decrease in animal speed during trajectories (Fig. 4A, C). Decoding error vs. spatial bin for both CA1 and PFC is shown in Fig. 4B (Kruskal-Wallis with Bonferroni post-hoc; CA1, Χ^2^(7) = 109.2, p = 1.2e-6 for spatial bin 5 vs. spatial bin 1; PFC, Χ^2^(7) = 44.19, p = 0.03 for spatial bin 4 vs. spatial bin 1), and animal speed in spatial sections during trajectories is shown in Fig. 4C (Kruskal-Wallis with Bonferroni *post-hoc*; Χ^2^(7) = 2165.4, p < 1e-100 for pairwise comparisons of spatial bins 3-6 to other bins indicating significant slowdown; p ≥ 0.16 for all spatial bin comparisons within bins 3-6). Running speed therefore had an effect on position representations in both CA1 and PFC.

We next asked whether the theta-cycle position representations of CA1 and PFC ensembles were correlated, which would suggest that spatial information is shared across the two networks. Recent studies have found similar coherent spatial representations in CA1 and visual cortical areas (Haggerty and Ji, 2015; Saleem et al., 2018). Similar to these studies, we therefore examined the theta cycle correlations between decoding errors for CA1 and PFC estimates, while controlling for actual location and speeds. In this approach, correlation between CA1 and PFC decoding errors is computed within each spatial section (spatial sections are used in order to control for the relationship between speed and location during shuffling), and then compared to correlation values with shuffled data while controlling for location and speed (see also **Materials and Methods**), i.e., shuffled correlation values for statistical comparison are obtained by shuffling CA1 and PFC data across speed-matched theta cycles within each spatial section.

We indeed found that CA1 and PFC decoding errors within theta cycles were correlated with each other (Fig. 4D-G). The distribution of CA1 vs PFC decoding error values were distributed along the diagonal with peak around 0, but were significantly correlated, similar to previously reports for CA1 and visual cortex (Saleem et al., 2018). Fig. 4D illustrates the distribution of raw correlation values for CA1 vs. PFC decoding errors for one spatial section, and average for all spatial sections in Fig. 4E; average correlation: r = 0.10, n = 10 sessions, two-tailed t-test of actual correlations compared to uniform distribution; t(9) = 4.60, p = 0.001). Crucially, in order to control for effects of speed modulation, we compared this actual correlation to values obtained from shuffled data, where CA1 or PFC decoding error was shuffled across time bins (theta cycles) within individual spatial sections, and within the same speed category (shuffled time bins had the same speed range - low, mid, or high, as the original time bins), thus preserving the relationship between speed and position (see also **Materials and Methods**, (Saleem et al., 2018)). Fig. 4F shows the difference between actual and shuffled correlations, with residual decoding errors distributed principally along the diagonal, indicating that CA1 and PFC ensembles had correlated spatial coding. Indeed, a comparison of actual vs. shuffled correlation values for all 10 behavior sessions (Fig. 4G) showed that actual decoding error correlations were significantly higher than shuffled values, with a decrease from 0.10 to 0.03 (paired-sample two-tailed t-test; n = 10; significant difference between actual and shuffled correlations, t(9) = 3.19, p = 0.01). These results therefore suggest shared and coherent coding of spatial position in the two regions governed by theta oscillations.

### Theta phase modulation of CA1 and PFC activity

In addition to coherence between theta oscillations in CA1 and PFC, spiking activity in also known to be modulated by theta phase in the two regions. Phase-locked spiking and phase precession in CA1 are well-established phenomena known to result in improved spatial coding by providing a temporal code for position owing to a relationship of spiking within place fields to the phase of ongoing theta oscillations itself (O’Keefe and Recce, 1993; Skaggs et al., 1996; Harris et al., 2002; Schmidt et al., 2009) (Fig. 5). Similarly, several studies have established prominent phase-locking of PFC spiking to hippocampal theta oscillations, and shown that this phase-locked spiking is correlated with performance in spatial memory tasks (Jones and Wilson, 2005a; Benchenane et al., 2010; Spellman et al., 2015). Phase precession in PFC neurons with respect to hippocampal theta oscillations has also been reported (Jones and Wilson, 2005b). Whether this theta-phase associated spiking in PFC results in improved spatial coding is not clear.

In order to address this question, we first quantified theta-phase associated spiking, namely phase-locking and phase-precession in simultaneously recorded CA1-PFC populations (see **Materials and Methods**, Fig. 5). Similar to previous reports by others and us (Jones and Wilson, 2005a; Siapas et al., 2005; Jadhav et al., 2016), CA1 and PFC neurons exhibited strong phase-locking, with a large fraction of neurons in both regions phase-locked to theta oscillations (Fig. 5A-C; Rayleigh z-test; p < 0.05 criterion; CA1 fraction significant = 83.5% or 213/255; PFC fraction significant = 51.4% or 57/111). Fig. 5A shows illustrative examples of significantly phase-locked CA1 and PFC neurons, along-with their spatial firing preferences on a linearized trajectory. Note that these cells exhibit both spatial selectivity as well as preferred theta phase spiking within their spatial firing fields. Spiking in the firing fields is therefore not uniformly distributed across phase, but rather occurs at particular phases of theta oscillations for phase-locked neurons. Fig. 5B shows distribution of phase-locking strengths for the two regions, and distribution of preferred peak phases for significantly phase-locked CA1 and PFC neurons is shown Fig. 5C. As expected, phase locking strength was significantly higher for CA1 compared with PFC neurons (phase-locking strength measured as the kappa concentration parameter of a circular Von-mises distribution; CA1 vs. PFC phase locked neurons n = 213 vs. 57; two sample Kolmogorov-Smirnov Test; p = 1.5e-08). Preferred phases for CA1 and PFC phase locked neurons were also different, similar to previous reports (Kuiper two-sample test; the two distributions for preferred phases of the phase locked neurons are significantly different, T(kuiper)=5025, p = 0.002).

We next characterized phase-precession properties of CA1 and PFC neurons. Phase precession of place cells, or the advancement of firing from early to late theta phases as an animal traverses a place field, is a major constituent of CA1 theta modulation (O’Keefe and Recce, 1993; Skaggs et al., 1996; Schmidt et al., 2009; Feng et al., 2015), and weak phase precession has also been reported in PFC neurons (Jones and Wilson, 2005b). We quantified phase precession using a previously reported method (Kempter et al., 2012) that maximizes the circular-linear correlation of spike location versus spike theta phase, using the peak firing fields of each neuron (see also **Materials and Methods**). Similar to previous reports, we found a large fraction of CA1 neurons exhibited significant phase-precession (Fig. 5D, E, 58.0% neurons, 148/255 with p < 0.05 criterion), with a much smaller fraction of significant neurons with phase precession in PFC (6.3%, 7/111 with p < 0.05 criterion). Illustrative examples of phase precession for CA1 and PFC are shown in Fig. 5D (CA1 neuron, ρ = −0.64, p = 1.7e-7; PFC neuron, ρ = −0.27, p = 0.02; note that the example phase-locked neurons in Fig. 5A did not exhibit significant phase precession, p > 0.63 for both neurons), and the distributions of the phase precession parameter, Rho (ρ) (the R-value equivalent for goodness-of-fit for linear circular relationships) are shown for all CA1 and PFC neurons in Fig. 5E. These distributions were significantly different as expected (two sample Kolmogorov-Smirnov Test; p = 2.7e-4).

These results confirm previous findings of strong theta phase-locking and weak theta phase precession in PFC neurons (Jones and Wilson, 2005b; Siapas et al., 2005; Jadhav et al., 2016). Interestingly, as previously suggested (Jones and Wilson, 2005a; Benchenane et al., 2010; Spellman et al., 2015), they also show that phase-specific spiking in PFC exhibits spatial preferences and therefore phase relationships to spatial location similar to CA1 (Fig. 5A, D), raising the possibility of a phase code in PFC similar to CA1.

### Incorporation of theta phase improves spatial decoding in both CA1 and PFC

Theta phase is known to underlie an additional temporal code in CA1 that provides more accurate spatial information than just the firing rates alone, and indeed incorporation of theta phase has been shown to significantly improve hippocampal spatial decoding (Jensen and Lisman, 2000). Since PFC neurons also exhibit theta-phase specific spatial firing within firing fields (Fig. 5), we next asked whether this phase relationship can lead to improved decoding accuracy in PFC populations by taking into account these phase-space relationships, in parallel with CA1 ensembles.

We used a strategy similar to a previous study (Jensen and Lisman, 2000) for incorporating theta phase in population decoding of spatial position in both CA1 and PFC, in order to assess and compare the impact on spatial representation in these regions (Fig. 6). Spatial firing field templates were sectioned into an increasing number of evenly spaced phase bins for decoding analyses, with each phase-bin based template taking into account theta phase in addition to the firing field of the neuron. Each neuron’s firing field therefore contributes multiple templates (equivalent to number of phase bins used for decoding), which are spatially restricted *only if* a phase-based code for spatial position exists for the original firing field. Fig. 6A illustrates for a CA1 neuron how 7 phase bins result in templates with spatially restricted firing due to phase precession in the firing field, with the new phase bins encoding sub-parts of the original firing field. Similarly, for the PFC neuron illustrated in Fig. 6B, phase-locked spiking results in phase-based spatial templates due to phase locking in the firing field, with only the preferred phase bins (indicated with arrows) exhibiting spatially localized firing. Note that for a uniform or random distribution of phases in the firing field (i.e., no phase-space relationship), multiple phase bin templates will not increase the spatial localization, but rather distribute the firing field randomly amongst the phase templates. Therefore, an important control for this analysis is generating a similar number of templates but with shuffled phase bins, which preserve the increased number of phase-based templates but randomize the phase-space relationship. Similarly, simply using smaller time windows without reference to phase will also not trivially increase the spatial information, since smaller time windows led to higher decoding error in both CA1 and PFC (Fig. 6C), presumably due to reduction in number of spikes (Kruskal-Wallis with Bonferroni post-hoc; CA1, Χ^2^(3) = 263.1, p = 3.7e-9 for theta vs. 50 ms; PFC, Χ^2^(3) = 214.87, p = 3.7e-9 for theta vs. 50 ms).

Using this method for generating phase-based templates for population decoding, we found a significant and consistent improvement in spatial decoding accuracy in CA1 with increase in number of phase bins, similar to previous findings, with asymptotic improvement in decoding error at ~6-8 phase bins, ((Jensen and Lisman, 2000), Fig. 6D). There was a ~2-fold median improvement in CA1 decoding accuracy when 7-phase bins were taken into account (median (IQR) decode error for 1 phase bin: 3.4cm (1.4 to 8.3 cm); median (IQR) decode error for 7 phase bins: 2.0cm (0.9 to 3.8 cm); Kruskal Wallis with Bonferroni post hoc, Χ^2^(7) = 1277.47, p ≤ 0.02 for all pairwise successive phase-bin comparisons for bins 1-5, pairwise successive phase-bin comparisons greater than 5, p = 1). Using a similar decoding approach for PFC ensembles, we also found a significant and strong increase in spatial decoding accuracy for PFC populations, with a ~4-fold median improvement in PFC decoding accuracy when 7-phase bins were taken into account (Fig. 6E; median (IQR) decode error for 1 phase bin: 15.7cm (3.7 to 46.9cm); median (IQR) decode error for 7 phase bins: 3.6cm (1.4 to 30.6cm); Kruskal Wallis with Bonferroni post hoc, Χ^2^(7) =1684.76, p ≤ 0.0013 for all successive pairwise comparisons for bins 1-7, pairwise phase-bin comparison between 7 and 8, p = 0.24). The median decoding accuracy for PFC ensembles with 7 theta phase bins was comparable to the original CA1 decoding accuracy obtained with just with firing rates (median (IQR); CA1 with 1 phase bin, 3.4 cm (1.4 to 8.3 cm), PFC with 7 phase bins, 3.6 cm (1.4 to 30.6 cm); CA1 accuracy at 1 phase bin was still significantly higher than PFC at 7 phase bins, Wilcoxon rank sum test for comparison between the two; z = −13.28, p = 3.0e-40).

Incorporation of theta phase thus improves spatial decoding accuracy in both CA1 and PFC, suggesting that spatial firing fields in both regions are co-modulated by theta phase. In order to control for the possibility that the increased decoding accuracy may simply arise from increased number of templates, we performed the same analysis with shuffled firing templates, by re-assigning spikes to a random theta phase bin before computing new phase-binned fields (encoding shuffle). This shuffling procedure thus removes any phase-space relationships by assigning spikes from firing fields to random theta phases, but preserves the original firing fields for the increased number of templates (for e.g., for 7 phase bins, each of the new 7 templates will have the same overall firing field as the original field for the neuron, but with random assignment of phase bins). With these shuffled templates, we observed no improvement in decoding accuracy with increase in phase bins for either CA1 or PFC populations (Fig. 6E; pairwise Wilcoxon rank sum for shuffled phase bins in CA1 2-8 vs. regular phase bins 2-8, p < 2.6e-208; **Fig. 6F;** Pairwise Wilcoxon rank sum for shuffled phase bins in PFC 2-8 vs. regular phase bins 2-8, p < 1e-99).

The improved position decoding accuracy can therefore cannot be explained simply as an increase in number of templates, or the sharpening of templates due to a trivial decrease in number of spiking events in smaller time windows. Finally, we also examined whether correlations in CA1 and PFC spatial coding were preserved when theta phase was incorporated in population decoding. Using a similar approach as above (Fig. 4F), we computed correlations between actual decoding errors and shuffled errors for each session, illustrated for decoding errors obtained using 7 theta phase bins (Fig. 6H). Similar to correlations based on just firing rate decoding, we found that actual decoding error correlations were significantly higher than shuffled values, with a decrease from 0.13 to 0.02 (Fig. 6H; paired-sample two-tailed t-test; n = 10; significant difference between actual and shuffled correlations, t(9) = 3.05, p = 0.009). Incorporation of theta phase thus improves spatial decoding accuracy in both CA1 and PFC.

## DISCUSSION

Our results establish a theta-phase mediated mechanism of temporal coordination for coherent coding of spatial position in hippocampal-prefrontal networks during memory-guided behavior. We report two major novel findings; first, we found that prefrontal population activity encodes animals’ current position on a theta-cycle timescale during memory-guided behavior, and this prefrontal coding of spatial position is coherent with hippocampal coding. Second, we found that theta phase-associated spiking significantly improved prefrontal representations of spatial position, simultaneously with improvement in CA1 spatial representations, while maintaining coherent coding.

The physiological mechanisms that underlie hippocampal-prefrontal interactions are of great interest. It is well-established that these regions have complementary roles in memory processes, and multiple direct and indirect anatomical pathways between the two regions can support communication necessary for memory-guided behavior (Cenquizca and Swanson, 2007; Vertes et al., 2007; Ito et al., 2015; Spellman et al., 2015; Hallock et al., 2016; Shin and Jadhav, 2016; Maisson et al., 2018). Further, inactivation studies that target these interactions have reported deficits in spatial memory (Floresco et al., 1997; Riedel et al., 1999; Wang and Cai, 2006, 2008; Churchwell et al., 2010; Maharjan et al., 2018).

Theta oscillation mediated interactions are the most prominently implicated physiological mechanism for long-range hippocampal-prefrontal communication in spatial memory paradigms. Theta oscillations in the two regions exhibit oscillatory coherence, and more than half of prefrontal neurons exhibit phase-locked spiking to hippocampal theta oscillations (Hyman et al., 2005; Jones and Wilson, 2005a; Siapas et al., 2005; Benchenane et al., 2010; Gordon, 2011; Spellman et al., 2015; Shin and Jadhav, 2016; Guise and Shapiro, 2017). Importantly, coherence and phase-locking are enhanced during spatial memory performance, (Jones and Wilson, 2005a; Benchenane et al., 2010; Hyman et al., 2010; Hallock et al., 2016; Guise and Shapiro, 2017), and impaired in parallel with cognitive deficits in genetic knockout models (Sigurdsson et al., 2010; Harris and Gordon, 2015). Theta-mediated hippocampal-prefrontal interactions therefore support spatial memory-guided behavior, and there is evidence that these interactions can support mnemonic representations, such as task-selective activity, choice specific responses, and memory encoding for spatial working memory (Jones and Wilson, 2005a; Benchenane et al., 2010; Hyman et al., 2011; Spellman et al., 2015; Guise and Shapiro, 2017). Despite this evidence, whether and how theta oscillations support shared processing of spatial information, the most fundamental feature represented by hippocampal place cells, has remained unexplored. Further, although theta is known to mediate a phase-based code for position in CA1, whether theta-phase also influences PFC spatial coding is not clear.

In this study, we therefore had two major goals – examine spatial coding relationships in CA1 and PFC ensembles during memory-guided behavior, and investigate the effect of theta phase on spatial coding. Addressing these questions requires simultaneous population recordings in the two regions and application of Bayesian decoding analyses, which we implemented in animals performing a W-track spatial alternation task that requires hippocampal-prefrontal interactions (Maharjan et al., 2018). We confirmed that PFC neurons exhibited spatially specific firing (Fujisawa et al., 2008; Ito et al., 2015; Jadhav et al., 2016; Mashhoori et al., 2018), but as expected, CA1 place cells exhibited significantly higher spatial specificity. We then used Bayesian decoding analyses to extract moment-by-moment spatial position from ensemble activity. We used theta cycles as time bins, since a) theta oscillations provide temporal windows for organization across hippocampal-cortical networks (Jensen and Lisman, 1996; Lisman, 2005; Lisman and Redish, 2009; Mizuseki et al., 2009; Lisman and Jensen, 2013), and b) since we wanted to investigate the effect of theta phase on spatial coding. For this firing rate based Bayesian decoding, using fixed time bin of lengths close to the average theta cycle (~125 ms) yielded similar decoding error (Fig. 6), which increased with smaller time windows, presumably due to decrease in relevant spiking information.

Using this firing rate based Bayesian decoding, we found that PFC ensembles also encoded spatial position within individual theta cycles, but with higher decoding errors than CA1 as expected from their lower spatial specificity. Crucially, simultaneous population decoding allowed us to examine correlations in cycle-by-cycle decoding error. We found that CA1-PFC position representations were coherent, with decoding errors from both regions significantly correlated with each other, even when controlling for effects of speed. This coherent coding indicates shared processing of spatial signals in the CA1-PFC network, which can be supported by multiple direct and indirect connections. In particular, direct connections from ventral CA1 to PFC, as well as indirect connections from PFC to CA1 via nucleus reuniens, have been implicated in altering task selective spatial representations between the regions (Ito et al., 2015; Spellman et al., 2015), suggesting that a similar mechanism may underlie coherent spatial coding. These interactions can therefore be bi-directional, although we note that theta-mediated interactions have been shown to exhibit a primarily CA1-leading-PFC directionality (Gordon, 2011; Jadhav et al., 2016). It is therefore possible that coherent PFC spatial representations are influenced by CA1 spatial activity, which can be investigated in future studies. Interestingly, the degree of CA1-PFC spatial coherence we observed was similar in magnitude to recently reported correlations for CA1 and primary visual areas (Haggerty and Ji, 2015; Saleem et al., 2018), raising the possibility of shared spatial representations in widespread networks.

We next examined the effect of theta-phase associated spiking on joint spatial representations in the two regions. CA1 place cells exhibits theta phase precession, where spike timing relative to theta phase conveys finer spatial information than place field firing alone (O’Keefe and Recce, 1993; Skaggs et al., 1996). Phase precession is also related to place cell theta sequences, which represent sequential activity within individual theta cycles than sweep through positions near the animal’s current position (Feng et al., 2015; Wikenheiser and Redish, 2015). This phase-based temporal coding results in significant improvement in estimation of position from CA1 populations (Jensen and Lisman, 2000). We therefore reasoned that PFC spatial representations may also be similarly influenced, since a) theta oscillations mediate strong hippocampal-prefrontal interactions during memory-guided behavior, and b) PFC neurons exhibit prominent hippocampal theta phase-associated spiking. We confirmed strong theta phase-locked spiking in both CA1 and PFC regions (Jones and Wilson, 2005a; Jadhav et al., 2016), whereas theta phase precession was prominent in CA1 (O’Keefe and Recce, 1993; Skaggs et al., 1996) and weak in PFC (Jones and Wilson, 2005b). Both CA1 and PFC neurons thus exhibited a strong relationship between theta phase and spatial firing, with a preference to spike at particular phases within their spatial firing fields, underlying a phase-space relationship (Fig. 5). We asked if using theta phase as an additional parameter in population decoding leads to an improvement in position estimation. We indeed found that both CA1 and PFC spatial representations showed significant improvement in spatial coding when theta phase was taken into account. This improvement in spatial accuracy was not seen with shuffled phase assignments, where we generated a similar number of templates, but with random assignment of phase, thus disrupting the phase-space relationship. Theta-phase relationships therefore enhanced spatial accuracy, and also maintained coherent CA1-PFC spatial coding, suggesting shared spatial processing. Finally, PFC representations showed a greater improvement in spatial accuracy with phase than CA1, which was likely a floor-effect resulting from the lower limit of the spatial resolution of our position tracking (~2 cm), and the fact that CA1 firing rate representations were already highly accurate.

This increase in spatial decoding accuracy is likely due to a phase specific segregation of spatially selective spikes from multiple neurons with similar phase preferences, leading to a sharpening of spatial information when decoded spikes are in the preferred phase bin. Interestingly, we observed significant improvement in decoding accuracy with the use of up to 6-8 phase-based templates for theta cycles. It has been previously argued that for CA1, this corresponds to the phenomenon of theta-gamma coupling, where optimal improvement in decoding accuracy corresponds to number of nested gamma (~40-80 Hz) cycles within theta cycles (Jensen and Lisman, 2000). Theta-gamma coupling is also known to be related to CA1 theta sequences (Colgin, 2011; Zheng et al., 2016), and theta-gamma coupling has also been recently reported in hippocampal-prefrontal networks (Tamura et al., 2017), raising the possibility that a similar phenomenon can influence PFC spatial representations.

Our results thus establish that hippocampal and prefrontal neural populations coherently encode position during spatial memory behavior, and further that theta-phase underlies a temporal mechanism that concurrently improves spatial representational accuracy in both regions. We hypothesize that this phenomenon can possibly extend beyond local coding of current position. Interestingly, spatial decoding accuracy showed a behavioral-dependent decrease near choice points in both CA1 and PFC. Hippocampal theta sequences are known to be prominent during these behavioral epochs and support non-local representations of behaviorally relevant trajectories or upcoming goals (Johnson and Redish; Gupta et al., 2012; Wikenheiser and Redish). Our results raise the possibility that a similar mechanism of theta-phase mediated coordination can support non-local coding in hippocampal-prefrontal networks. By elucidating a physiological mechanism for coordination of spatial representations, these findings therefore provide an important foundation for future investigations of how theta oscillations organize shared mnemonic representations in hippocampal-prefrontal networks.

## Acknowledgements

This work was supported by NIH Grants R01 MH112661, a Sloan Research Fellowship in Neuroscience (Alfred P. Sloan Foundation), and Whitehall Foundation award to SPJ. We would like to thank the late John Lisman for inspiring the investigation of theta phase and prefrontal spatial representations.

